# Rhythmidia: a modern tool for circadian period analysis of filamentous fungi

**DOI:** 10.1101/2024.05.15.594281

**Authors:** Alex T. Keeley, Jeffrey M. Lotthammer, Jacqueline F. Pelham

## Abstract

Circadian rhythms are ubiquitous across the kingdoms of life and serve important roles in regulating physiology and behavior at many levels. These rhythms occur in ∼24-hour cycles and are driven by a core molecular oscillator. Circadian timekeeping enables organisms to anticipate daily changes by timing their growth and internal processes. *Neurospora crassa* is a model organism with a long history in circadian biology, having conserved eukaryotic clock properties and observable circadian phenotypes. A core approach for measuring circadian function in Neurospora is to follow daily oscillations in the direction of growth and spore formation along a thin glass tube (race tube). While leveraging robust phenotypic readouts is useful, interpreting the outputs of large-scale race tube experiments by hand can be time-consuming and prone to human error. To provide the field with an efficient tool for analyzing race tubes, we present Rhythmidia, a graphical user interface (GUI) tool written in Python for calculating circadian periods and growth rates of Neurospora. Rhythmidia is open source, has been benchmarked against the current state-of-the-art, and is easily accessible on GitHub.

## Introduction

From unicellular bacteria to multicellular eukaryotes, circadian rhythms are important regulators of organismal processes across the tree of life [1–3]. Circadian rhythms are biological processes that occur in a ∼24-hour oscillation. These rhythms allow organisms to accurately keep time and anticipate, rather than react to, daily environmental changes. This anticipatory behavior and physiological coordination can facilitate biological processes like growth, metabolism, and sleep at appropriate times of day [1,4]. In fungi, circadian rhythms time processes like spore development and release, allowing organisms to compensate against rhythmic damaging environmental effects [5]. For example, it may be advantageous to release thinner-walled, less UV-resistant spores at night for their protection, while more hydrophobic spores might be released during the drier daytime [5]. In the filamentous fungi *Neurospora crassa*, asexual spore production, termed conidiation, occurs predawn during high humidity and low temperatures, which are ideal conditions for spore dispersal [6].

Neurospora is a longstanding model organism for studying circadian biology, genetics, and cell biology due to its rich history and tractability for study. The Neurospora research community employs a plethora of well-established molecular biology and biochemical tools for investigation, including a fully sequenced genome and complete knock-out collection [7–11]. Eukaryotic circadian clock properties are also strongly conserved in Neurospora at both the micro- and macroscopic levels [11–13]. Using Neurospora, researchers first identified the central tenet of eukaryotic timekeeping, the negative feedback loop, which was subsequently identified as the primary timekeeping mechanism in animals [11,14]. Furthermore, the nature of light and temperature entrainment of the clock and the mechanism of clock output through control of gene expression was first studied and conceptualized in Neurospora [11,15,16].

Circadian rhythms are subject to entrainment, the process by which an organism’s biological clock is synchronized with external environmental cues such as light, temperature, or nutrients to align to the proper phase of the oscillating stimuli [1,17]. However, without entrainment cues, an organism’s internal circadian oscillator will continue to cycle under constant conditions (e.g. continuous darkness). In constant conditions, the length of the endogenously generated period by the central oscillator is known as the “free-running” period, which for Neurospora is about 22.5 hrs [18]. With this in mind, if the circadian period could be accurately and reliably measured, one could decode the molecular basis for clock function by assessing how genetic or environmental changes alter the free-running period when Neurospora is grown in continuous darkness.

To better understand the mechanics of the circadian oscillator and its regulation of physiology, it is essential to measure and observe circadian phenotypes. One aspect of Neurospora growth that is under control of the circadian clock is spore production. Specifically, the clock times the production of conidia, asexual spores. Conidiation is achieved when vegetative mycelia differentiate into aerial hyphae that produce macroconidia (Figure 1a) [10,19]. A standard tool to examine Neurospora phenotypic rhythms of conidiation is the race tube assay (Figure 1a). The assay involves using race tubes, which are ∼40 cm long glass tubes bent upward at the ends and filled with a layer of agar growth media (Figure 1a). Neurospora conidia are inoculated at one end of the race tube (Figure 1a). After inoculation, race tubes are typically grown in constant light (LL) for ∼24 hours and then transferred to constant darkness (DD) to synchronize the rhythms. The absence of light allows for growth under constant conditions and calculation of the free-running circadian period of the strain. Every day, the growth front is marked on each tube to provide a temporal reference frame for analysis. After circadian culturing is complete, the tubes are then photographed from the bottom using a flatbed scanner or similar instrument to capture and convert growth into digital information (Figure 1b).

**Fig. 1.**
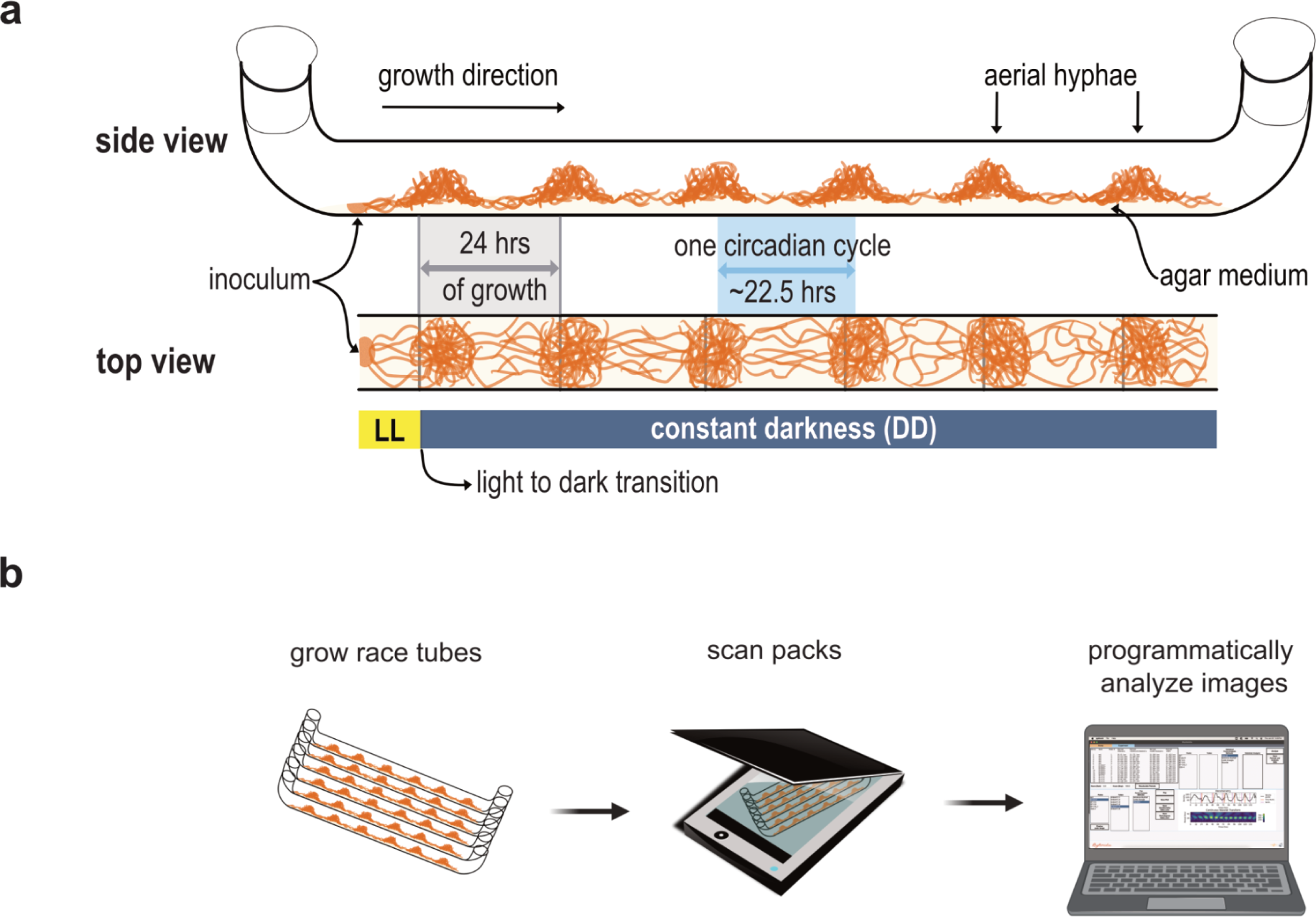
Race tube anatomy and experimental workflow. **a.** A schematic of a race tube assay. Race tubes are filled with ∼15mL of agar medium and inoculated with conidia at one end. The tubes are then incubated in constant light (LL) for 24 hrs, followed by a transition to constant darkness (DD). The Neurospora continues to grow with the clock controlling development and morphology at the growth front. For a specific portion of the relative circadian day, the mycelia differentiate into aerial hyphae and produce asexual spores (conidia, orange meshwork, referred to as banding). The peak-to-peak distance of conidial banding corresponds to one relative circadian cycle (highlighted in blue, ∼22.5 hrs) and the free-running period of the strain. To establish a relationship between growth and time, it is necessary to mark the tubes once every ∼24 hrs (vertical gray lines) at the growth front. Time marks serve as representative time frames for circadian period calculation. **b.** High-level workflow of race tube assays. Neurospora is grown as described above in packs of six race tubes over approximately a week. Once growth reaches the ends of the tubes, packs are scanned from the bottom using a high-resolution flatbed document scanner. From this point, race tube images can be cropped or otherwise preprocessed before being uploaded to Rhythmidia for programmatic analysis.

The daily marking of the tubes establishes a linear relationship between time and distance within the race tube, assuming a sufficiently constant growth rate throughout the experiment (Figure 1a). By measuring the distances between sites of conidial banding, it is possible to calculate the average circadian period of a strain. Implementing this manual method of period calculation from race tubes has several drawbacks: performing dozens of measurements and calculations by hand for multiple packs of six tubes per experiment is time-consuming and prone to human errors. Additionally, manual measurements can yield inconsistencies in determining at what point in a conidial band to conduct the measurements. Furthermore, areas of increased conidiation are broad, and the exact locations of peaks of conidial density are only sometimes apparent to the human eye. While digitization and computational solutions mitigate many of these complications, no recent advances have been made since 2008 [20]. The first image analysis software for race tube analysis, Chrono II, was developed in Pascal starting in 1990, by Roenneberg and Taylor [20]. This software’s successor, ChronOSX, is the current standard and state-of-the-art for analysis of race tube experiment images.

While ChronOSX has been the workhorse for race tube analysis since its initial release in 2000, it has some limitations. First, ChronOSX is closed-source software, impeding customization and contributions from the broader Neurospora community should additional functionality be desired. Second, ChronOSX is not freely available online. Third, ChronOSX only runs on older Macintosh operating systems. The most recent operating system ChronOSX will function on is macOS Sierra, originally released in 2016. Finally, ChronOSX has not been updated since 2008, despite major advances in computational image processing since that time. Considering these factors, we feel the field is overdue for advancement.

Here, we introduce Rhythmidia, a graphical user interface (GUI) for Python version 3.11 and above, which facilitates the calculation of circadian periods from race tube assays. Rhythmidia guides the user through programmatic upload and orientation of scanned race tube images and user-mediated algorithmic identification of individual race tubes, time marks, and conidial banding present in the image. Furthermore, Rhythmidia allows users to save and revisit analysis data and perform four different methods of calculating circadian periods from analyzed images. Rhythmidia can be installed and run as a freely available Python package, with the code and documentation hosted on GitHub. Running in Python means that Rythmidia will continue functioning on any modern computer running a compatible version, requiring minimal time and effort to install and run.

### Design and Implementation

Rhythmidia is a GUI-based Python package for analyzing circadian periods and growth rates in Neurospora race tube assays. Given the user base Rhythmidia is aimed at and the visual process of race tube image analysis, a GUI-based implementation is essential. The general experimental and analytical workflow starts with inoculating and growing one or multiple packs of six race tubes (Figure 1b). After culturing, these packs are scanned to generate race tube images. These images are then uploaded to the Rhythmidia user interface (Figure 2), and the software facilitates user-mediated identification of image features before saving and analyzing the data. Users can view data for each uploaded race tube image in an experiment file, including the periods calculated by four different methods. Furthermore, Rhythmidia performs basic statistical analysis of any combination of saved data on a by-pack or by-tube basis. The experimental data tab also allows users to visualize densitometry and periodogram plots and export any experimental or analytical data (Figure 3).

**Fig. 2.**
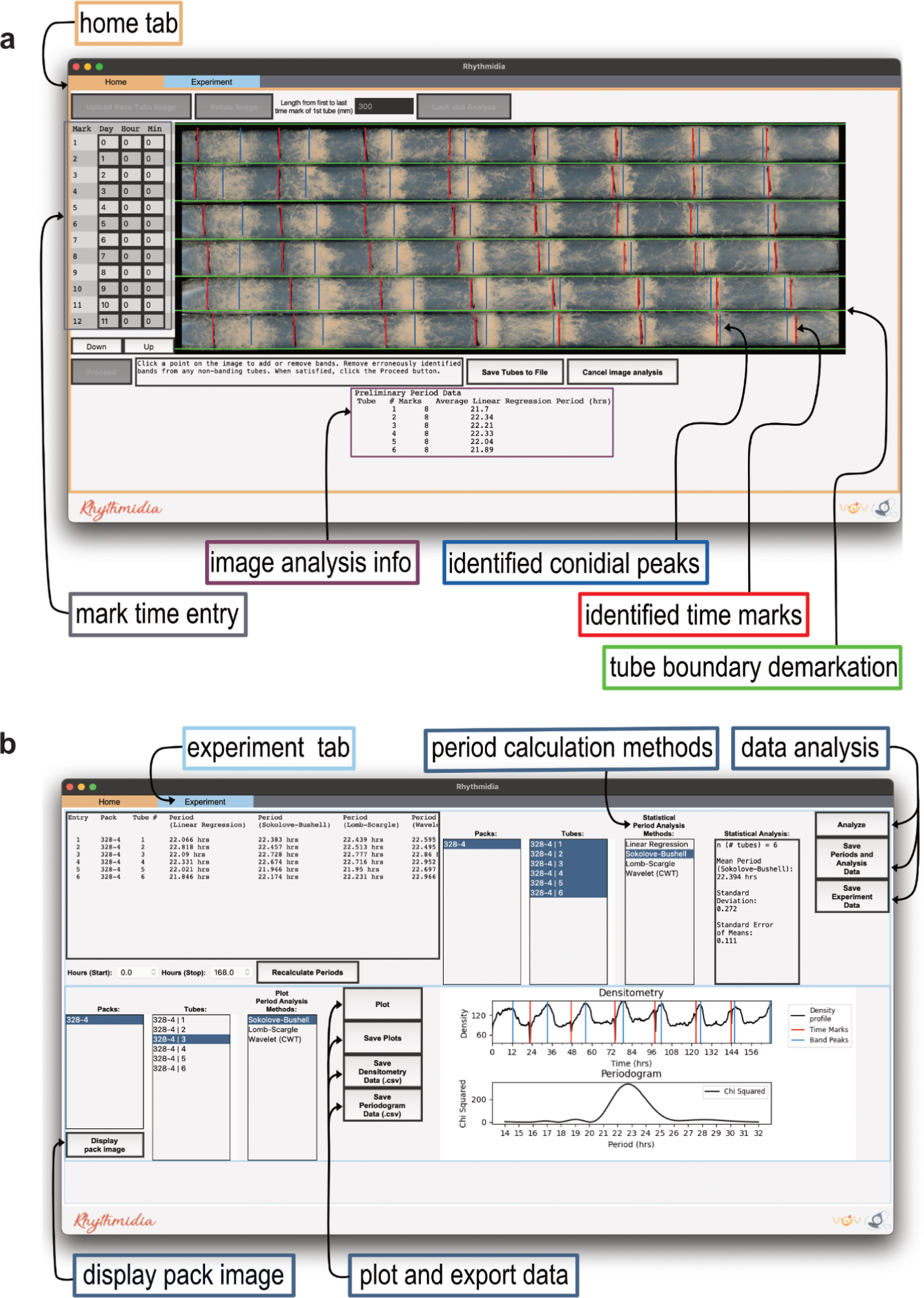
Rhythmidia interface. **a.** Rhythmidia image analysis interface of the home tab with programmatically identified features of a race tube image highlighting tube boundary demarcations (green), time marks (red), and conidial banding peaks (blue) overlaid on an uploaded image. An entry field for tube mark times is located to the left of the image (grey), and preliminary linear regression periods are displayed below the image (image analysis info, purple). The top of the screen holds “tabs” for switching between the image analysis (home tab) and the experimental data analysis interfaces. **b.** Rhythmidia experiment and data analysis interface, displaying data from a single six-pack of race tubes. Includes per-tube calculated periods and growth rate, options for statistical analysis of multiple tubes’ and packs’ periods, and options for plotting densitometry and period analysis data of individual tubes, all by multiple period analysis methods. The interface also includes options to export plots, data, and statistical analysis output.

**Fig. 3.**
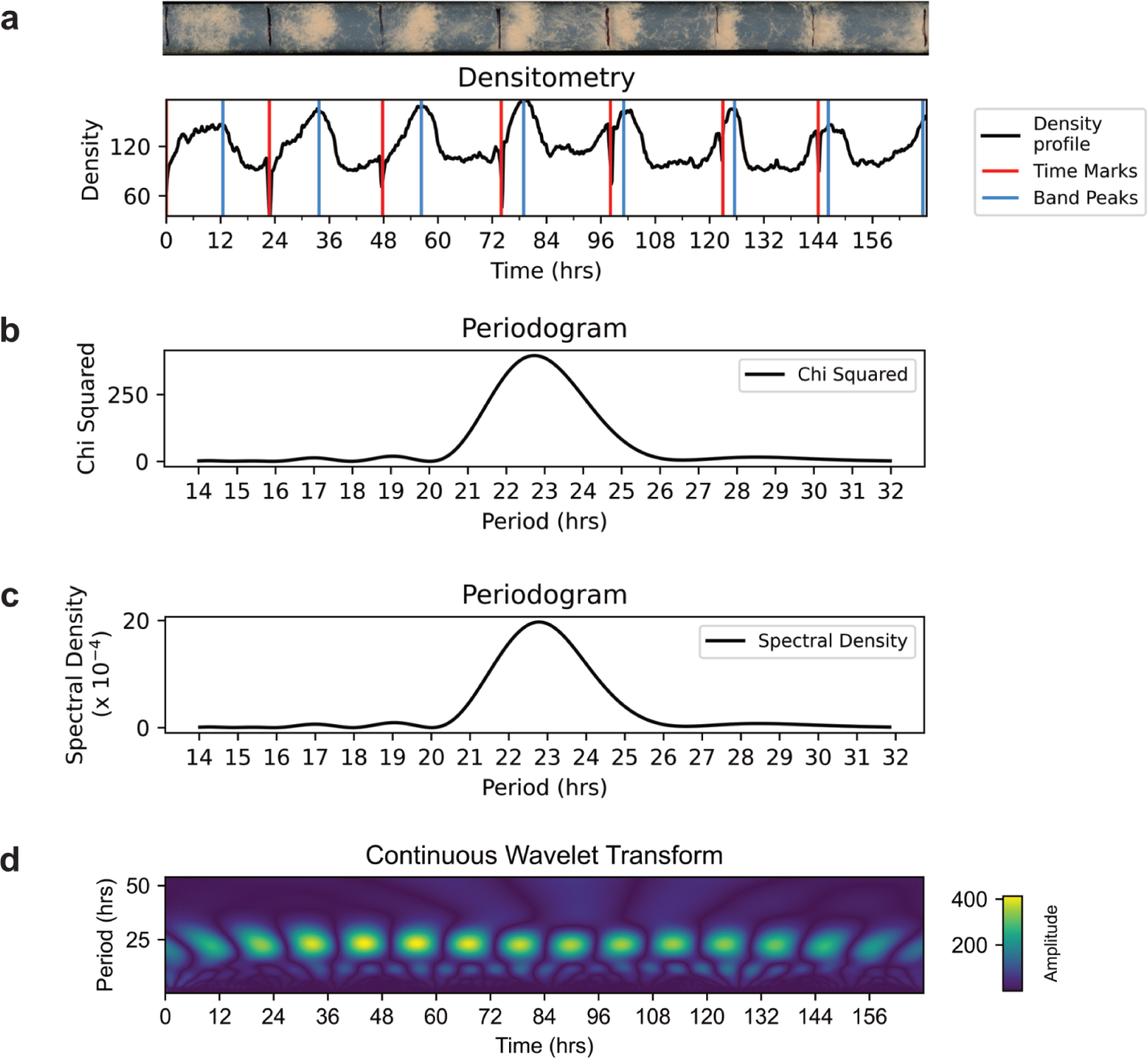
Rhythmidia exports. **a.** Example plotted raw densitometry output from corresponding race tube above. Each of the three output period analysis plots from Rhythmidia, the (**b.**) Sokolove-Bushell periodogram, (**c.**) Lomb-Scargle periodogram, and (**d.**) continuous wavelet transform heatmap.

Rhythmidia accepts images in various image formats (S1 Table), and can analyze images as-is in terms of color and contrast. Once imported, all images are automatically internally converted to greyscale and resized to 1160×400 pixels upon upload using bicubic interpolation rescaling (S1 Fig). In a greyscale image format, each pixel of an image is designated a single number between 0 and 255, conveying the color, or “brightness,” of the pixel numerically from black (0) to white (255). Handling images in such a format allows Rhythmidia to normalize the colors of different race tube images to a single data dimension per pixel. An image size of 1160×400 pixels reflects the aspect ratio typical to the straight portion of a pack of six race tubes, and any stretching that may occur in a particular image does not impact downstream calculations more than the error of the measurement (S2 Fig). Furthermore, 1160×400 pixels is the maximum optimal size for GUI display on modern 13-inch screens and larger within the context of the Rhythmidia analysis interface.

### Canny edge detection and probabilistic Hough transforms for tube boundary identification

After the image is successfully uploaded, resized, and converted to greyscale, Rhythmidia begins image feature detection (S2 Fig). The first algorithm Rhythmidia uses to identify image features involves the detection of Canny edges in uploaded images. Canny edge detection is a computer vision algorithm for identifying edges within images [21]. The algorithm relies on a Gaussian filter to smooth out noise in an image before identifying the intensity gradient of the image. Thresholds are then applied to the magnitudes of the gradient to identify edges in the image. Using edges identified by Canny or similar algorithms, a progressive probabilistic Hough transform can be used to identify straight lines of a minimum length in an image [22].

In the case of Rhythmidia, this minimum length threshold is set such that any sufficiently long lines would have to correspond to boundaries between race tubes. In order to identify the likely margins demarcating the boundaries between individual race tubes in uploaded images, Rhythmidia first uses the scikit-image package’s module for Canny edge detection [23], with low and high thresholds set to 1 and 25, respectively. Rhythmidia then performs a progressive probabilistic Hough transform of these edges using the scikit-image implementation to identify a likely set of sufficiently long horizontal lines of sufficiently low slope present in the image (S1 Fig).

### Vertical densitometry for tube boundary demarcation

Depending on the image, noise from scan quality or growth of the organism may cause the Canny edge method of horizontal line detection to be insufficient. To circumvent this issue, Rhythmidia employs a second method to improve edge detection involving vertical density profiles of the image (S1 Fig). Averages of pixel brightness are taken along horizontal line segments at each vertical pixel of the image, both at the left edge of the image and in the center (S1 Fig, red traces). Rythmidia then identifies places where local minima exist in each density profile with sufficiently close y values corresponding to a line of low slope passing through both points. Here, it draws lines of demarcation between race tubes that pass through the centers of the pair of sampling regions (S1 Fig, blue lines).

Edge detection using Canny edges and the Hough transform to identify lines separating race tubes in the image is effective in cases where areas of low conidial density in images are distinct from the image’s background. Rhythmidia uses the vertical density profile model to identify horizontal lines for cases where this distinction is unclear. In every case, the software performs both the Canny edges method and the vertical density method of horizontal line detection and selects the larger set of acceptable lines to present. This gives the highest chance of correctly identifying the actual boundaries between race tubes present in the image. These lines demarcating the boundaries between individual race tubes are extended to the image’s width and subject to user review (Figure 2a, green lines).

### Noise filtering and feature detection via densitometry

Before Rhythmidia can perform circadian analysis, it must identify and discriminate between both time marks and conidial banding patterns in the race tube images (Figures 1a and 2a). Rhythmidia achieves this by creating a one-dimensional density profile of the brightness of each identified race tube, referred to as the raw density profile (S1 Fig). Rhythmidia then performs a filtering process on these density profiles to more efficiently identify features within each race tube to reduce noise (S3 Fig). Rhythmidia employs the SciPy Savitzky-Golay filter to successively fit polynomial functions to the raw density profile of each tube using the linear least squares method [24]. To improve the capacity for conidial band identification, a fourth-order polynomial is used to smooth the raw density profile (S3 Fig) [24,25]. Next, to help identify time pen marks in the raw densitometry plot, Rhythmidia uses the Savitzky-Golay filter with a window size of 30 pixels by fitting an eighth-order polynomial (S3 Fig). In each case, the lowest possible order polynomial is applied to preserve the characteristics of peaks in the density profile.

After smoothing the noise, Rhythmidia determines the time marks in each race tube in the image by identifying extreme local minima in the density profile, corresponding with the dark lines created by the pen marks used to keep experimental time (S4 Fig). Time mark minima are broadly identified by the SciPy peak finding algorithm using two criteria. The first criterion for selection is whether the minimum is of sufficient prominence from the base of the peak, and the second is if the minimum has a minimally steep flanking slope; if these two criteria are met, the respective location is selected as a likely time mark (S4 Fig, black arrows) [24]. The time marks manifest as technical artifacts that appear in the density profile as extreme local minima. Time mark artifacts are problematic as they would be interpreted as low levels of conidial density; thus, correcting for these extreme local minima before performing period calculations is essential.

Rhythmidia corrects for time mark artifacts by modifying the density profile by excluding the identified pen marks from the data by interpolating density near these pen marks’ positions (S4 Fig). The interpolation is accomplished by generating data between the density values flanking the excluded regions. Simply stated, it connects the “minima gap” in the data, an artifact of the time mark, with a straight line between the data points on either side of this gap (S4 Fig).

Next, Rhythmidia identifies likely positions of conidial banding within each race tube by using the filtered and interpolated density profile to identify local maxima with sufficient width and prominence. These maxima are likely locations of maximal conidial banding (S4 Fig, blue arrows). Any falsely identified time marks or improper identification of conidial peaks can be immediately rectified by the subsequent user-mediated quality control step, in which users can simply click to add or remove presumptive time marks before any calculations occur (Figure 2a, red and blue lines). The interpolated densitometry data is used to identify conidial banding sites and for densitometry analysis for circadian period calculation.

### Circadian period analysis

Once race tube boundaries, time marks, and conidial band peaks are identified within the uploaded image, information is saved to a file (termed the experiment file) for each race tube identified in the image. This includes a user-specified name for the pack of tubes in the image, the identification number of the tube in the image (numerically from the top down), the name of the original image file, and the greyscale version of the original image. It also includes tube-specific information: the y boundaries of the tube within the image, the x positions of the pen marks and conidial banding sites within the tube and corresponding times, the calculated mean growth rate of the tube, and the raw density profile of the tube.

The data from the analyzed image are then used to calculate the circadian periods of the strains in each tube and pack. The associated statistics are displayed in the “Experiments” tab (Figure 2b). Rhythmidia employs four different methods for circadian rhythm analysis: linear regression period calculation, the Sokolove-Bushell periodogram, the Lomb-Scargle periodogram, and the continuous wavelet transform [26–33]. A pair of dropdown selection menus in the experiment interface allows users to perform these analyses on a region of the experiment between any two marked time points.

#### Linear regression period calculations

The linear regression method of calculating circadian periods involves multiplying the distance between conidial banding sites by the inverse of the growth rate per hour for the given tube. These values are calculated by measuring the distances between pen marks and dividing these distances by the differences between the corresponding times of marking during the race tube experiment (Figure 1a). Rhythmidia achieves this by calculating a mean rate of growth in pixels per hour using the positions of time marks within a tube and user-provided time data and a mean distance in pixels between conidial banding sites within the same tube.

#### Sokolove-Bushell periodogram calculation

In addition to linear regression period calculations, Rhythmidia uses the Sokolove-Bushell periodogram to calculate circadian periods of race tube images. The Sokolove-Bushell periodogram is a valuable tool for investigating circadian rhythms within biological systems. The method, rooted in statistical principles, involves the application of the chi-square statistic to assess periodicities in time-series data, which adapts the chi-square test for non-uniformly sampled data and is particularly well-suited for studying biological phenomena characterized by irregular temporal sampling [27,34,35]. Since its introduction in 1978, the Sokolove-Bushell periodogram has been widely used in circadian rhythm analysis and cited in over 1,000 publications, including in human, mammalian, ichthyic, bacterial, and fungal systems [20,36–38]. The approach involves binning the data into time intervals and calculating the chi-square statistic for each frequency, thus providing a robust measure of rhythmicity [27,34]. This method enables researchers to identify and quantify circadian patterns in biological processes, contributing to a nuanced understanding of temporal organization. The Sokolove-Bushell chi-squared periodogram for a given period *p* is equal to Q_p_;

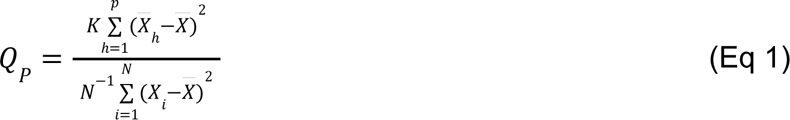

where *X̅_h_* is the mean of a set *K* values of length *P* under each time unit of the length of period *p*, and *X̅* is the mean of all N values of the data set over the time series [27].

In order to calculate circadian periods using the Sokolove-Bushell periodogram, Rhythmidia uses the corresponding function provided by the SciPy package for Python to generate a periodogram based on the density profile of a given race tube [24]. Rhythmidia then converts the output, provided in frequencies, to periods in pixels and then to hours using the mean conversion rate of pixels to hours calculated from the pen mark locations in the given race tube image and the user-provided time data. This periodogram then contains the power spectrum of each potential period of the density profile, with a peak corresponding to the period of best fit (Figure 3b).

#### Lomb-Scargle periodogram calculation

The Lomb-Scargle periodogram, rooted in the foundational works of Lomb (1976) and Scargle (1982), is an invaluable analytical tool in biology for unraveling temporal patterns within unevenly sampled time-series data, overcoming limitations associated with traditional Fourier-based techniques [28]. This method involves fitting sinusoidal functions at different frequencies to the observed data. By employing a least-squares optimization process, the periodogram quantifies the strength of periodic components across various frequencies, enabling the identification of rhythmic patterns in biological processes [28,29]. In brief, for a time course of N_t_ measurements X_j_ at t_j_, shifted so the mean is 0, the Lomb-Scargle periodogram at frequency *f* is equal to P_n_(f);

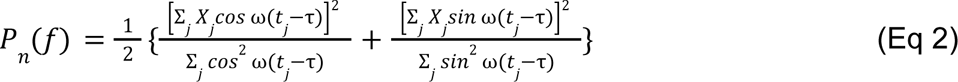

where ω ≡ 2π*f* [28,29,39].

In order to calculate circadian periods using the Lomb-Scargle periodogram, Rhythmidia uses the corresponding function provided by SciPy to generate a periodogram based on the interpolated density profile of a given race tube [24]. Rhythmidia then converts the output to periods in pixels from the original image. These can then be converted from pixels to hours using the pen mark locations in the given race tube image and the user-provided time data. This periodogram then contains the power spectrum of each potential period of the density profile, with a peak corresponding to the period of best fit (Figure 3c).

#### Wavelet analysis

The final period analysis method Rhythmidia employs is the continuous wavelet transform. The linear regression method and Fourier-based periodograms operate under the assumption of a single period throughout a time series and can thus miscalculate a single period for a series with a changing period or identify a series as aperiodic [40]. The continuous wavelet transform (CWT), on the other hand, computes a convolution of a time series that correlates each data point with a chosen wavelet function to determine the period at each point in time [30,32]. Wavelet functions can be stretched and translated to deconvolve even transient, high-frequency periodic signals [41,42]. Moreover, the continuous wavelet transform has a history of being used to identify changing or transient periods in circadian data [40,43,44].

Rhythmidia uses the PyWavelets package for Python developed by Lee *et al.* to apply the continuous wavelet transform to smoothed and interpolated race tube densitometry data and calculate periods over the time series in uploaded images [45]. PyWavelets takes in time series data and outputs a three-dimensional matrix for which the axes are time, frequency, and the amplitude of the wavelet transform. Rhythmidia takes this output and converts the frequency values to periods in hours, enabling the creation of a heatmap and the identification of the period of maximal amplitude and, thus, best fit at each time point of the experiment (Figure 3d). In order to do this, Rhythmidia identifies maximal amplitudes for each time point and accepts any greater than 75% of the maximum amplitude of the plot. The period for the tube is calculated as the mean of all accepted maximal amplitudes, and the continuous wavelet transform slope is calculated from the line of best-fit passing through all accepted maximal amplitudes.

#### Statistical analysis of calculated periods

Rhythmidia provides the user with an interface to perform statistical analysis of a subset of any periods calculated by any of the four listed methods for any race tubes saved to the same experiment file (Figure 2b). For any subset of periods selected, the mean, standard deviation, and standard error are computed using NumPy’s corresponding modules [46].

#### Growth rate calculation from image features

Analysis of fungal growth rate is a phenotypic screening method that connects to several biological processes, including the circadian clock. Tracking fungal growth rates using the glass tube method has been implemented since 1925 and popularized in Neurospora research by Beadle and Tatum in the 1940s [47,48]. It was one of the critical techniques that led them to their one gene-one enzyme hypothesis, garnering Beadle and Tatum the Nobel Prize in Physiology in 1958 [49,50].

Rhythmida can calculate growth rates via the user-provided measurement between the first and last time marks in a race tube, establishing a relationship between real distance and pixels. It uses the measured distance between the race tube’s initial and final time marks to establish a relationship between millimeters and pixels. Then, the mean growth rate of each tube is calculated using the elapsed times and distances between time marks. This involves the same linear regression method used for period calculation, resulting in race tube growth rates in millimeters per hour. Although not required, if the user inputs the measurement in the home tab, the calculated growth rate will appear in the experiment tab in the field that lists entries, packs, tubes, etc. If no measurement is provided, the experiment tab will read not applicable (“N/A”) for the growth rate column.

## Results

### Evaluating Rhythmida performance

To evaluate Rhythmidia in its methods of circadian period calculation, we benchmarked against the only comparable analysis tool available, ChronOSX [20]. To benchmark and compare Rhythmidia and ChronOSX, we analyzed the periods of six race tubes using the two overlapping methods present in both programs (linear regression (LR) and Sokolove-Bushell (SB) periodograms (Figure 4a). We grew a six-pack of strain 328-4, a classical banding control, and analyzed the same race tube image with Rhythmidia and ChronOSX. We found agreement in the mean calculated periods for Rhythmidia and CronosOSX of calculated periods between 22.2 and 22.7 hours respectively (Figure 4a).

**Fig. 4:**
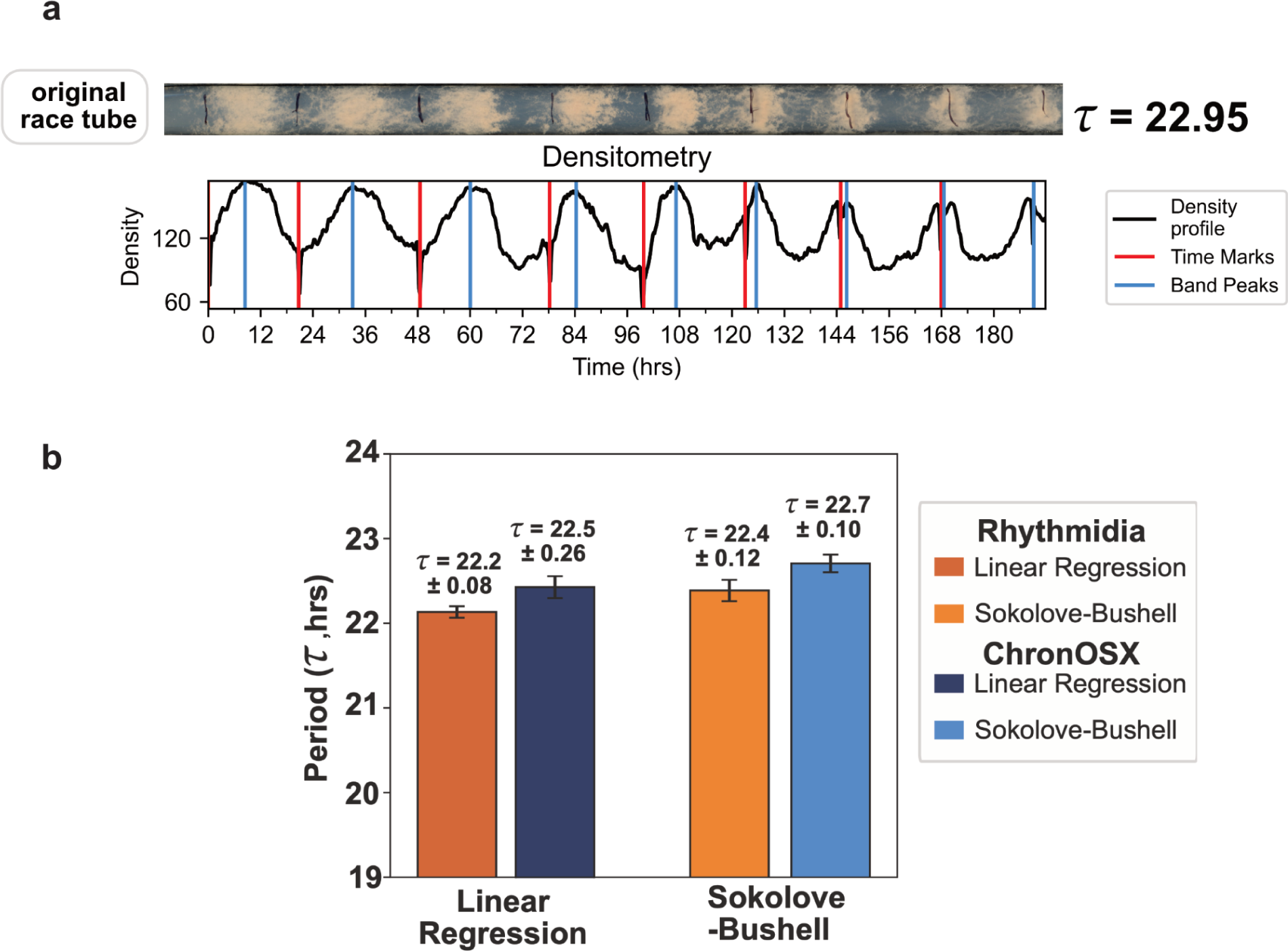
Rhythmidia performance. **a.** Example FGS race tube (at 5.55mM Sorbose) demonstrating 9 days of banding and subsequent densitometry profile. The period was calculated using the linear regression method. **b.** Comparison of periods calculated for the same pack of race tubes by Rhythmidia and ChronOSX, using linear regression (LR) and the Sokolove-Bushell (SB) periodogram. The bar plot reports the average period (*τ*) calculated by the reported method (n=6) and ± indicates the standard error of the means (SEM). The image used for analysis is in Supplemental figure 1a.

### Rhythmidia can handle longer data collection windows

ChronOSX can only analyze a maximum of seven days of data from race tubes. Depending on the growth rate of the strain and the precise length of the race tube, an experiment can yield 10-12 days of data or more; therefore, some data is subsequently excluded from ChronOSX analysis. To circumvent this issue, we built Rhythmdia with no limit to the number of possible days of analysis. To test the practical experimental limits of the Rhythmidia analysis window, we employed a strategy to slow the growth rate while preserving circadian function and capturing more bands and days of data.

Sorbose slows and restricts Neurospora growth [51–53]. Sorbose-containing media has been extensively used to restrict or slow the growth of Neurospora in various applications. It is most commonly used during electroporation for the genetic transformation of Neurospora and is referred to as Fructose-Glucose-Sorbose (FGS) or Sorbose-Fructose-Glucose media. The F:G:S ratio is typically 40:1:1, and is typically used at 0.1M sorbose for growth restriction to the colony size. To increase the number of banding days by slowing the rate of growth, we replaced the sugar content of our race tube media with FGS. We prepared race tube media containing FGS at 5.55 mM and inoculated with strain 328-4. This concentration yielded 9 days of banding, and using Rhythmidia’s extended data calculation capacity yielded a period of 22.95 hrs (Figure 4b).

## Discussion and Conclusion

While banding race tube experiments have been conducted since William Brandt first published the presence of the Neurospora banding phenotype in 1953, circadian period calculation came later with the work of Pittendrigh and colleagues in 1959 [18,54]. Modern analysis of circadian rhythms in Neurospora with race tube assays calls for an accurate, efficient, and repeatable approach for large-scale experiments. Current technological and experimental standards can leverage powerful computing and image analysis techniques to build upon decades of proven analyses. Furthermore, software that is easy to use for all computational skill levels while accurately employing field standard analytical methods for circadian data is critically missing in the field. Therefore, we sought to equip the research community with the ability to perform efficient, accurate analysis of race tube assays with an accessible open-source option: Rhythmida.

We developed Rhythmidia to improve the current state of the art in all of these regards, and did so with accessibility in mind. Rhythmidia has an easy-to-navigate user interface (Figure 2), allowing users to re-open files containing previous experimental data and analysis *ex post facto*. Rhythmidia also accommodates color accessibility, allowing users to select colors for image analysis and data plotting by hex code or color wheel. Moreover, Rhythmidia is more efficient than the current state of the art in allowing for efficient data export and plotting, enabling users to collate with their preferences (Figure 3). The benchmarks we applied to Rhythmidia demonstrated periods consistent with field literature for classical banding control strains, with results and errors comparable to the existing state-of-the-art, ChronOSX (Figure 4a) [18,55,56]. Furthermore, we developed Rhythmidia to analyze larger sets of data and tested this ability using growth-restricting reagent FGS (Figure 4).

## Availability and Future Directions

Rhythmidia is an open-source Python package. Its fully accessible and installable code is available on GitHub at https://github.com/Pelham-Lab/rhythmidia, complete with clear documentation and installation instructions. Rhythmidia is free to use under the MIT license. Moreover, as an open-source package written in Python 3.11 we anticipate extending new functionality and/or accommodating contributions from the broader Neurospora community through code contributions, bug reports, and feature requests. In particular, we plan to include methods for analyzing luciferase time course experiments as well as other experimental modalities.

## Experimental Methods

### Neurospora strains and culture conditions for race tube assays

The Neurospora strain used for race tube images was 328-4 mat A, bd^+^. Assays were performed in race tubes available from Chemglass Life Sciences (CG-4020-10) containing 15 mL of media (1x Vogel’s salts, 1.5% agar, 0.1% glucose, 0.17% arginine, 50ng/mL biotin). FGS experiments were conducted by replacing the 0.1% glucose with the corresponding concentration of FGS. In all cases, race tubes were poured and allowed to set overnight to decrease condensation. Tubes were inoculated with between 5.6×10^5^ (non-FGS media) and 4×10^6^ (FGS media) conidia (strain 328-4) suspended in 10μL sterile water from 5-7 day old minimal slants. Post-inoculation race tubes were incubated in constant light (LL) at 25°C for 24 hours, then transferred to constant darkness (DD) at 25°C until growth reached the far end of the tube. Each tube was marked every 24 hours at the growth front. Tubes were scanned using an Epson GT-1500 at 1200 dpi. Images were generated as .jpg and were uploaded to and analyzed using ChronOSX on a Macbook Pro running Mac OS X Leopard (10.5.8), and Rhythmidia on a Macbook Air M1 running MacOS Sonoma (14.1.1).

## Acknowledgments

We thank the Hurley Lab for gifted Neurospora strains. We thank Till Roenneberg for his insight on ChronOSX. We thank Jennifer Hurley and Alex Holehouse for their manuscript suggestions. Funding for this work was supported by the WashU Department of Biochemistry and Molecular Biophysics Cori Fellowship to J.F.P. J.M.L. was supported by the National Science Foundation via grant number DGE-2139839.

## Author Contributions

**Conceptualization:** ATK JFP

**Data curation:** ATK JFP

**Formal analysis:** ATK

**Funding acquisition:** JFP

**Investigation:** ATK JFP

**Methodology:** ATK JML JFP

**Project administration** JFP

**Resources:** JFP

**Software:** ATK JML JFP

**Supervision:** JFP

**Validation:** ATK JML JFP

**Writing – original draft:** ATK JFP

**Writing – review & editing:** ATK JML JFP

