## Supplementary material for "Rhythmidia: a modern tool for circadian period analysis of filamentous fungi": SI Keeley et al Rhythmidia sub1.pdf

### Supporting information

**S1 Fig**

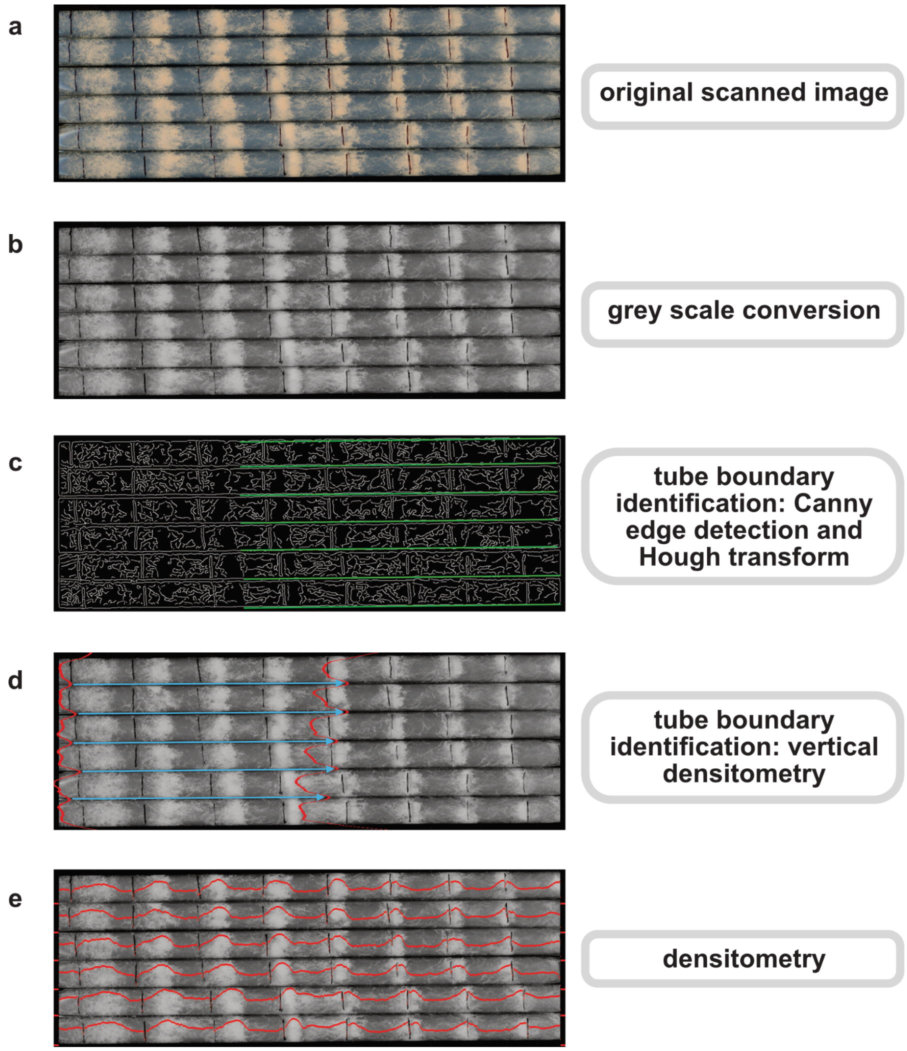


**S1 Fig: Algorithmic image feature identification: a.** Original scan of 6-pack of representative race tubes of strain 328-4 to depict algorithmic feature identification. **b.** Representative 6-pack post-Rhythmidia greyscale conversion. **c.** Depiction of detected Canny edges in representative race tube image to identify lines demarcating individual race tubes (green lines). **d.** Vertical bilateral densitometry (red traces) of a race tube image for identification of lines demarcating individual race tubes. Blue arrows indicate horizontal tube demarcation lines drawn between corresponding density minima. **e.** Raw densitometry of each race tube overlaid upon the corresponding race tube image.

### S2 Fig

a

b

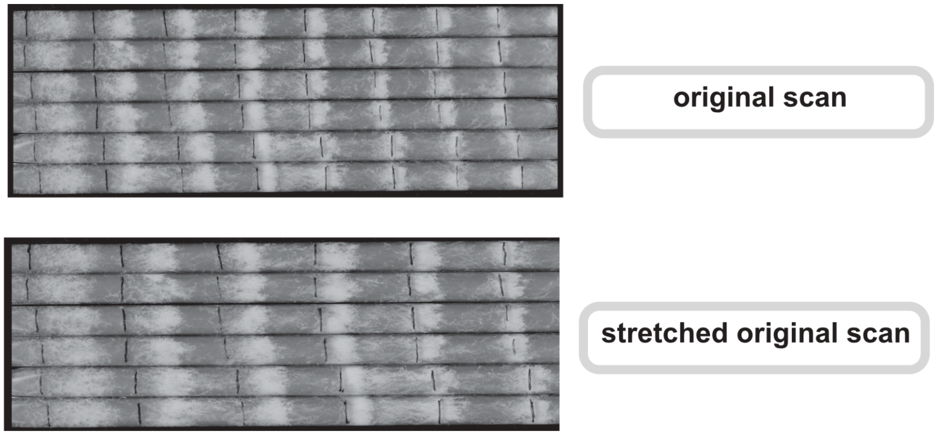


| Strain & Image | Period ($\tau$, hrs) | SEM | Period ($\tau$, hrs) | SEM | Period ($\tau$, hrs) | SEM | Period ($\tau$, hrs) | SEM |
|---|---|---|---|---|---|---|---|---|
| | **Linear Regression** | | **Sokolove-Bushell** | | **Lomb-Scargle** | | **Continuous Wavelet Transform** | |
| **328-4** | 22.29 | 0.23 | 22.23 | 0.14 | 22.31 | 0.12 | 22.31 | 0.09 |
| **328-4 Stretched** | 22.14 | 0.15 | 22.20 | 0.14 | 22.30 | 0.12 | 22.25 | 0.08 |

**S2 Fig: Image stretching period comparison.** Rhythmidia greyscale image output (**a.**) of analyzed image, as well as of cropped version of same image after programmatic resizing (**b.**, stretching). The able below displays calculated periods ($\tau$) in hours for both versions of the image, calculated using the linear regression, Sokolove-Bushell periodogram, Lomb-Scargle periodogram, and continuous wavelet transform. For each method, the period is calculated for each of six tubes, and these calculations are used to calculate the arithmetic mean and the standard error of the mean (SEM) for n=6 tubes.

### S3 Fig

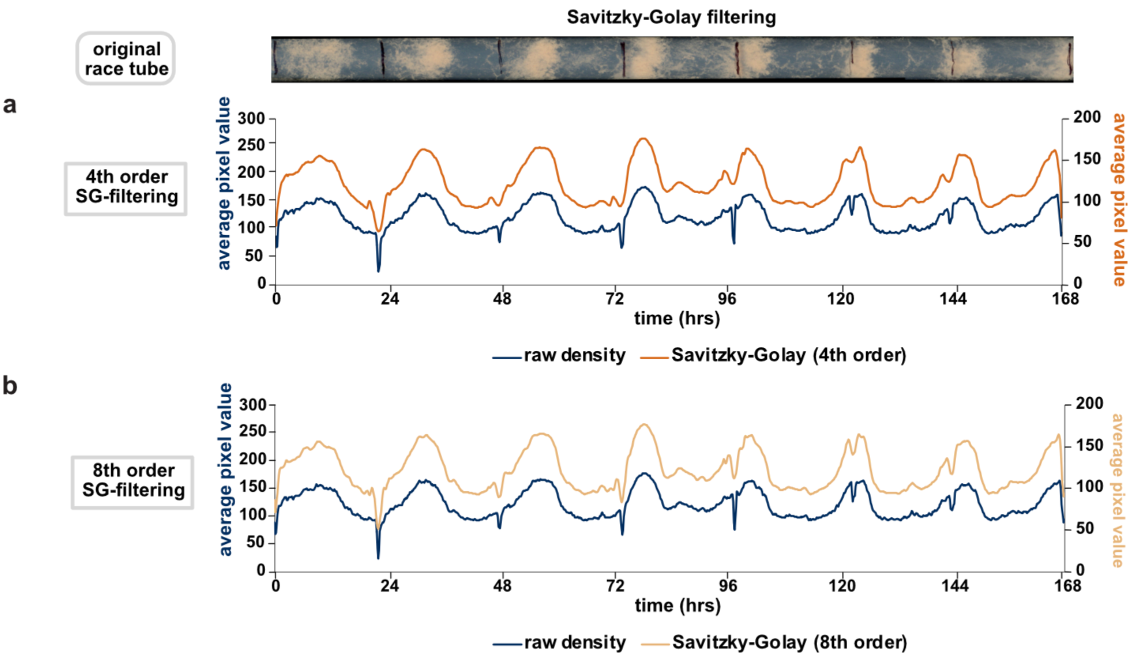


**S3 Fig: Savitzky-Golay filtering.** A single race tube image, aligned with representations of the effects of Savitzky-Golay filtering. Each plot compares the filtered to the raw density profile of a race tube using fourth-order (**a.**) and eighth-order (**b.**) fitting. Density is assessed as an average pixel value, with higher values corresponding to brighter pixels in a greyscale image.

**S4 Fig**

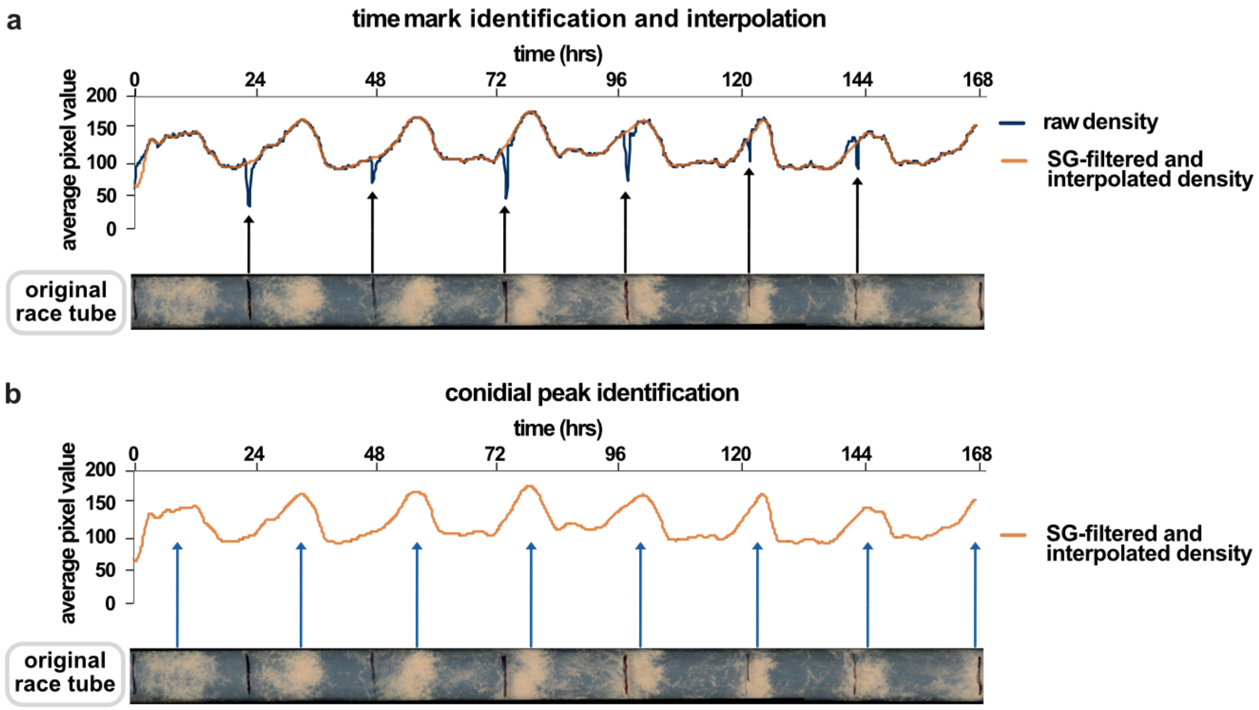


**S4 Fig: Time mark interpolation and conidial peak identification. a.** Comparison of raw density profile overlaid with smoothed, interpolated density profile of a single race tube, aligned with the original image of the tube. Black arrows connect time marks in the image to corresponding local minima artifacts in the raw density profile. **b.** Smoothed, interpolated density profile of a single race tube, aligned with the original image of the tube. Blue arrows indicate sites of conidial banding (peaks) in the image to corresponding local maxima in the density profile.

| File Type | |
|---|---|
| **.png** | Portable Network Graphics |
| **.tif and .tiff** | Tagged Image File Format |
| **.jpg and .jpeg** | Joint Photographic Experts Group |
| **.svg** | Scalable Vector Graphics |

**S1 Table:** List of accepted image formats of race tube images for upload to Rhythmidia's user interface.

### S1 Text: Rhythmidia installation and tutorial

*Installation*

Rhythmidia must be installed in a Python environment running Python 3.11 or higher using pip. Rhythmidia runs on Unix-like operating systems including macOS and Linux.
In order to install Rhythmidia, simply run the command:

```
pip install rhythmidia
```

This will install Rhythmidia and all of its dependencies. After installation, Rhythmidia can be launched from the command line by running the command:

```
rhythmidia
```

The terminal session used to run Rhythmidia must remain active while the software is in use.

*First-Time Use*

The first time you open a new installation of the program, you will be prompted to select a working directory. From this directory, the program will, by default, look for race tube images, and will by default, save data. This can be changed later at any time. You are not restricted to using this directory, it is purely for your convenience. This is also where analysis data will be exported by default.

Usage note: On some newer Mac laptops with variable-force-click trackpads, you may find that some clicks are not picked up effectively unless you click with full force.

*Image Preprocessing*

1. Crop your scanned race tube image as desired in the image viewer program of choice, leaving a small amount of background on either long edge

2. No need to make your image greyscale, fix its rotation, or increase contrast- Rhythmidia will take care of all of this internally

3. Race tube images can be .png, .tif, .tiff, .jpg, .jpeg, or .svg

*The Home Tab: Uploading and Analyzing an Image*

1. Upon opening the software, you will be greeted with the "Home" tab, which will look like this:

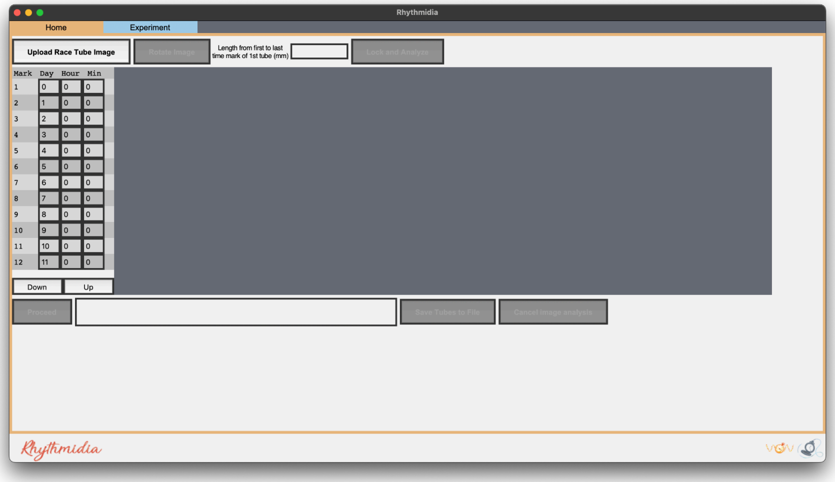


   a. NOTE:If you want to add tubes to an existing experiment file, go to File -> Open experiment File (or press ⌘O), although this is not necessary

2. To upload a new race tube image, click the button labeled "Upload Race Tube Image"

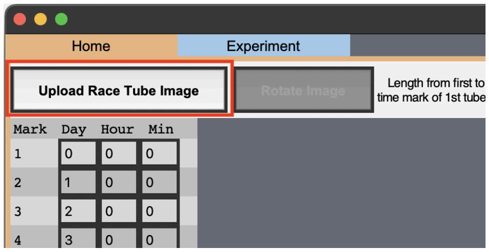


   a. Your image will appear in the center of the screen

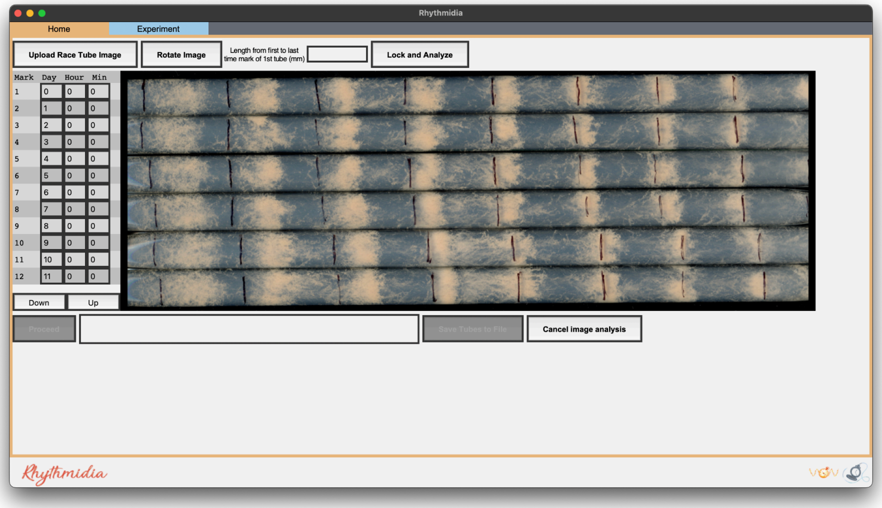


b. Uploading an image enables the options to rotate and to lock & analyze your image

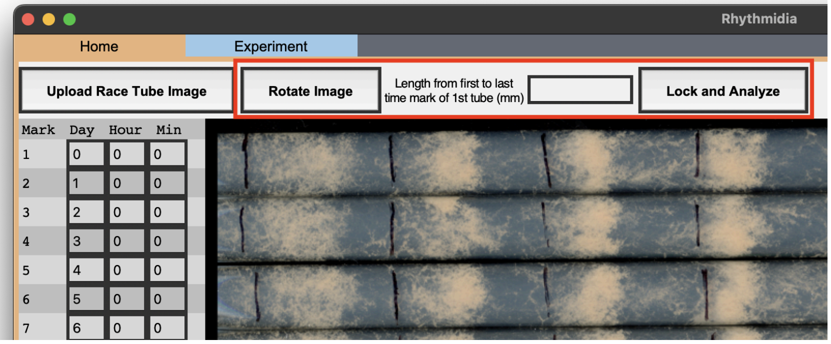


3. Rotate your image so that the growth direction of the tubes is from left to right across your screen
    a. To rotate your image 90 degrees clockwise, click the button labeled “Rotate”

4. When you are satisfied with your image’s orientation, click the button labeled “Lock & Analyze Image”

5. Rhythmidia will try to identify horizontal lines corresponding to the horizontal boundaries of the tubes in your image, including the lower and upper bounds below and above the final and first tubes
    a. One line between each two tubes

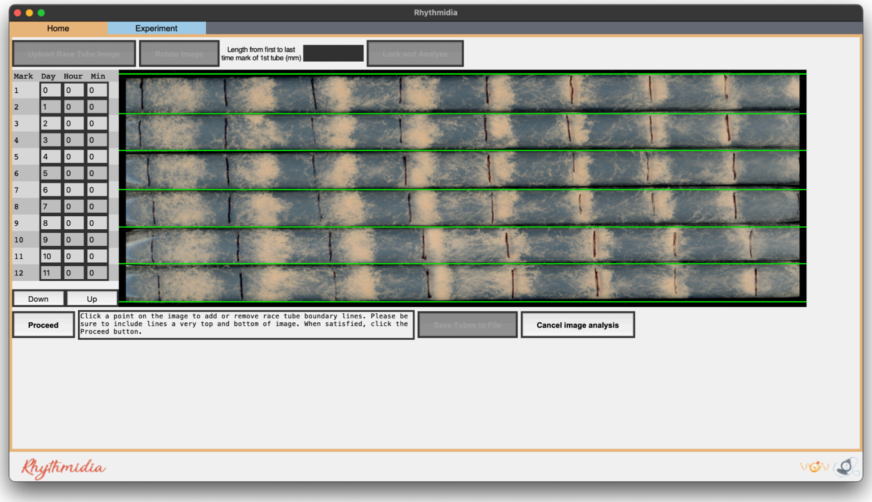


6. You will be directed to verify these lines:
    a. To remove an incorrect line, simply click on the line
    b. To add a missing line, simply click in an unoccupied position on the image

7. When you are satisfied with the positions of all tube demarcation lines, click the button labeled “Proceed”

8. Repeat steps 6-8 for time marks (red) and for the middle or peak of the bands (orange)

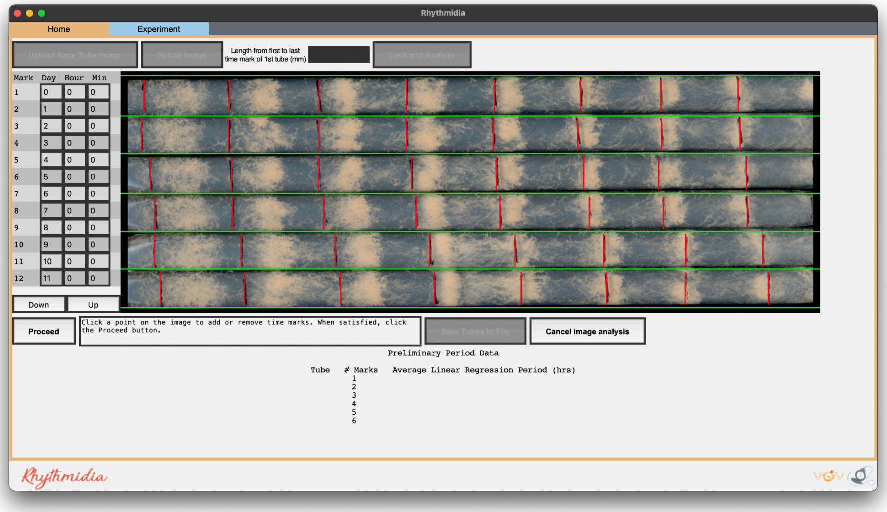

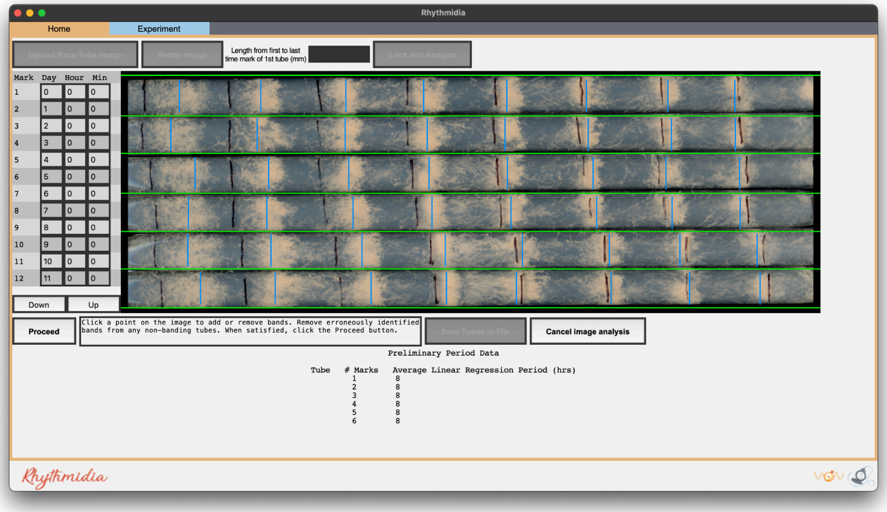


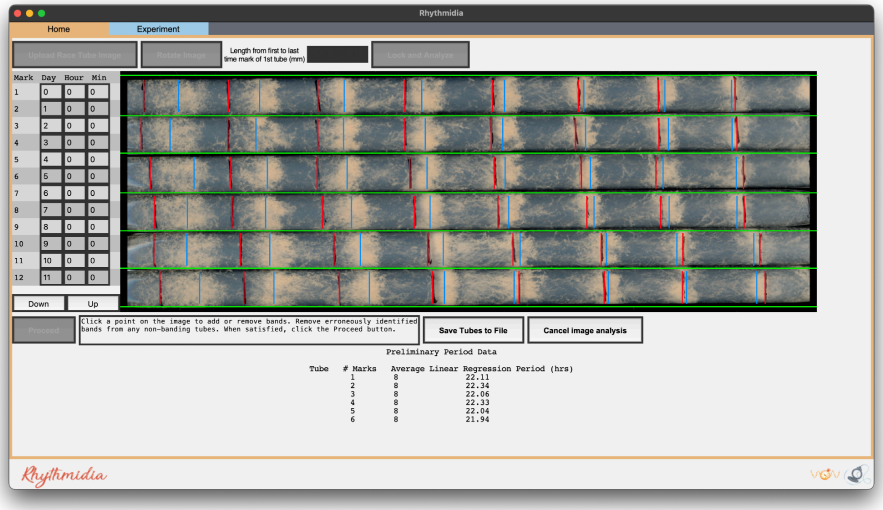


a. NOTE: At any time before saving tubes to the file, you may click the button labeled “Cancel image analysis”, which will reset the image analysis process and remove your uploaded image, while leaving open any open experiment file
b. NOTE: Be certain to record any differences in marking times in the mark sheet (left) before proceeding further. If tubes were marked at the same time every day, leave as the default setting (0 for all)
c. NOTE: The time marks will temporarily disappear while marking conidial peaks

9. After you are satisfied with the positions of the bands and click “Proceed”, you will be able to see a preliminary calculation of the period of each tube below

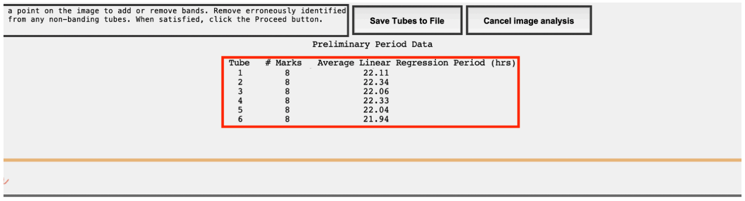


a. NOTE: if there is an issue at this stage (i.e. a missed or duplicated identifier) cancel image analysis and reload the image

10. You will now have the option to click the button labeled “Save Tubes to File”
    a. This will bring up a popup asking for a name for the pack of tubes in the current image before it saves them to file

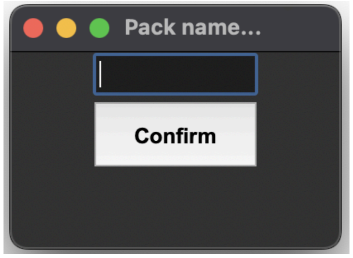


    b. If you are working within an existing experiment file, this will simply add this pack to the file and update it
    c. Otherwise, you will be prompted to Save As a new experiment file for these tubes
    d. Click on the Experiment tab to proceed

*The Experiment Tab*

1. Whether opening an existing experiment file or working from a new pack image, granular experiment data, plots, and statistical analysis data are located on the Experiment tab

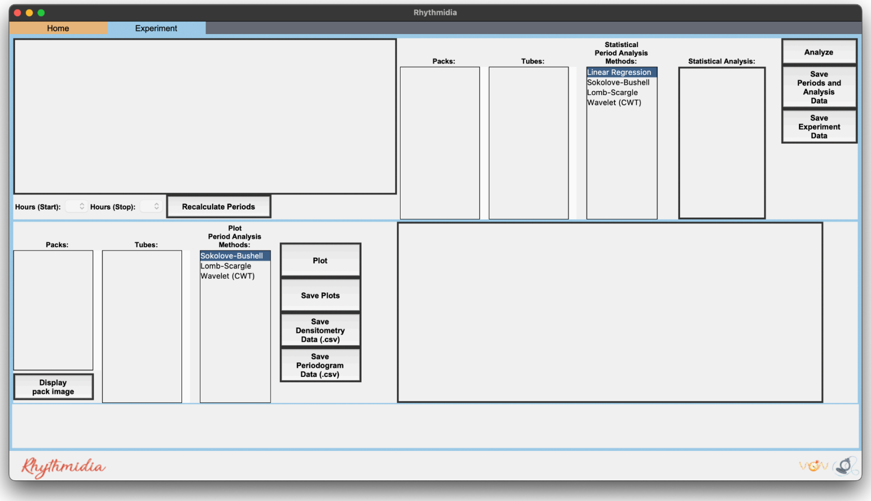


2. Experiment data (Entry, Pack, Tube # in pack from top to bottom, Period calculations, and Growth rate) is located in the table in the top left

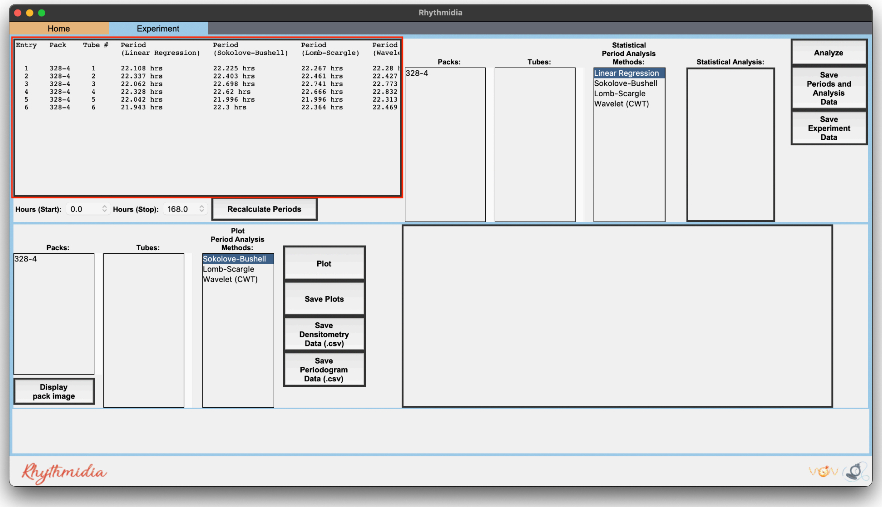

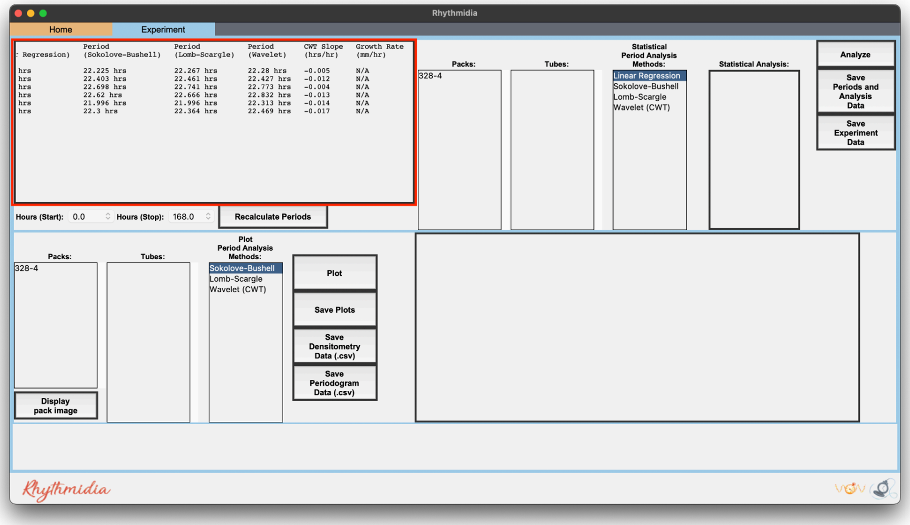

3. In the top right is the frame for statistical analysis of any number of tubes:

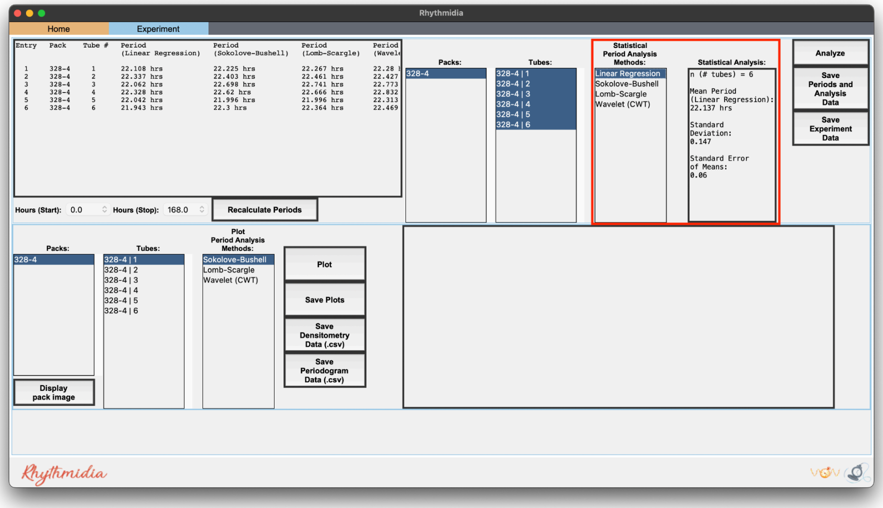

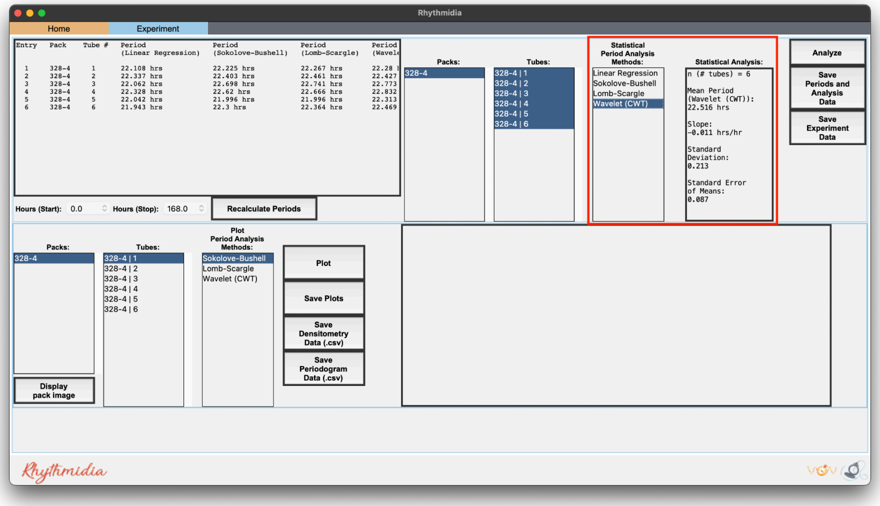

a. Select packs, tubes, and a method of period analysis in the 3 lists
b. To select multiple packs or tubes, use control-click
c. Click the button labeled “Analyze” to generate mean period, standard deviation, and standard error
d. Click the button labeled “Export Data” to export a .csv of the data for each tube selected
e. Click the button labeled “Export Analysis” to export a .csv of the analysis of the selected tubes

4. In the bottom half is the plot frame for plotting densitometry and a periodogram/heatmap of a single tube:
    a. Select pack, tube, and type of periodogram in the 3 lists
    b. Click the button labeled “Plot” to generate a densitometry plot and periodogram of the selected data
    c. Click the button labeled “Save Plot” to save an image of the dual plot in file format of choice

d. Click the button labeled “Save Densitometry” to save a .csv of the densitometry data
e. Click the button labeled “Save Periodogrammetry” to save a .csv of the periodogrammetry data

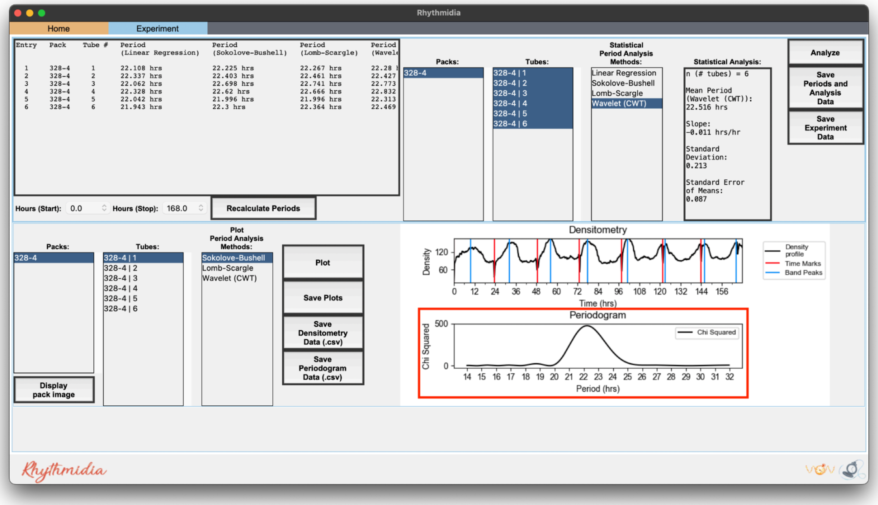


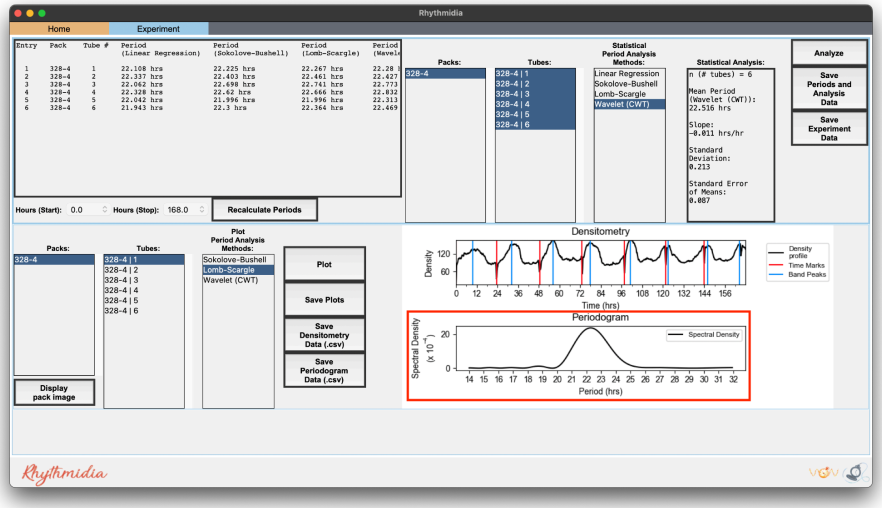

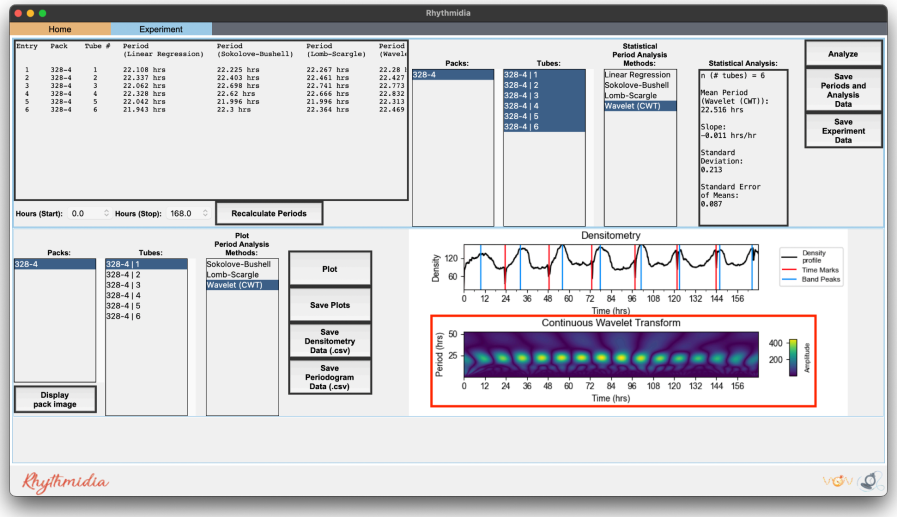


5. At the bottom left is a button labeled "Display Pack Image":
    a. this button will display a popup window containing the greyscale version of the image corresponding to whichever pack is selected in the bottom left list that was the exact image used for analysis

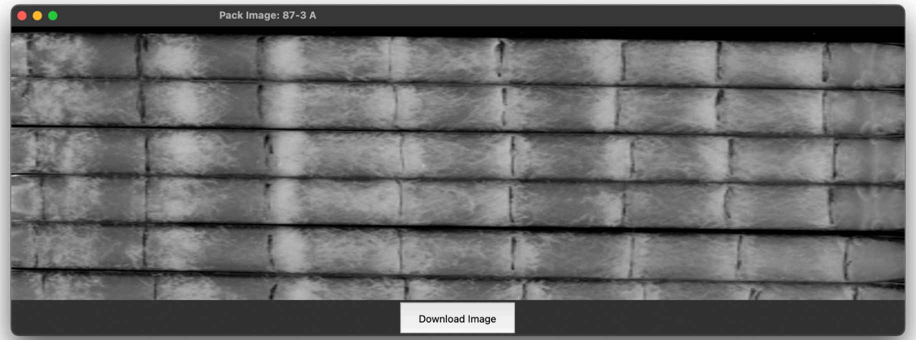
